# Functional characterization of sensory neuron membrane proteins (SNMPs)

**DOI:** 10.1101/262154

**Authors:** Hui-Jie Zhang, Wei Xu, Quan-mei Chen, Le-Na Sun, Alisha Anderson, Qing-You Xia, Alexie Papanicolaou

## Abstract

Sensory neuron membrane proteins (SNMPs) play a critical role in the insect olfactory system but there is a deficit of functional studies beyond *Drosophila*. Here, we provide functional characterisation of insect SNMPs through the use of bioinformatics, genome curation, transcriptome data analysis, phylogeny, expression profiling, and RNAi gene knockdown techniques. We curated 81 genes from 35 insect species and identified a novel lepidopteran SNMP gene family, SNMP3. Phylogenetic analysis shows that lepidopteran SNMP3, but not the previously annotated lepidopteran SNMP2, is the true homologue of the dipteran SNMP2. Digital expression, microarray and qPCR analyses show that the lepidopteran SNMP1 is specifically expressed in adult antennae. SNMP2 is widely expressed in multiple tissues while SNMP3 is specifically expressed in the larval midgut. Microarray analysis suggest SNMP3 may be involved in the silkworm immunity response to virus and bacterial infections. We functionally characterised SNMP1 in the silkworm using RNAi and behavioural assays. Our results suggested that *Bombyx mori* SNMP1 is a functional orthologue of the *Drosophila melanogaster* SNMP1 and plays a critical role in pheromone detection. Split-ubiquitin yeast hybridization study shows that BmorSNMP1 has a protein-protein interaction with the BmorOR1 pheromone receptor, and the BmorOrco co-receptor. Concluding, we propose a novel molecular model in which BmorOrco, BmorSNMP1 and BmorOR1 form a heteromer in the detection of the silkworm sex pheromone bombykol.

## Introduction

With the increasing throughput of draft genome sequencing, we are now in a position to complete a genome project within months. These genome projects are however working drafts and may harbor misassemblies or misannotations, requiring substantial curation before making biological inferences. To date there has been limited appreciation for professional curation by the wider genomic community. By curation here, we not only refer to a manual edition of the underlying gene model (i.e. structural annotation) but also functional assignment. This is an important activity: Any automated assignment of orthology – such as one based solely on the information contained within the protein sequences – does not rely upon, or imply, an orthology of function. Further, with incomplete ascertainment or without a robust phylogenetic framework, an assignment of orthology by descent may also be erroneous.

It is generally a challenge to identify accurately structural or functional annotations for diverged genes. An additional tool that underpins our ability to produce higher quality of both structural and functional annotations is expression profiling. Using RNA-Seq (i.e. high throughput sequencing), we can accurately detect both the open reading frames and untranslated regions of transcripts. In addition, expression level profiling RNA-Seq and quantitative PCR (qPCR) can provide strong evidence for functional annotation assignment. Indeed, together with a knock-down/out technique such as RNA interference (RNAi) or Clustered Regularly Interspaced Short Palindromic Repeats (CRISPR), these approaches give us one of the strongest lines of evidence what the function of a gene may be. Having accurate assignments is important to the wider biology community: The traditional concept of nomenclature implies that genes between two species that have the same name are orthologous by either function or descent.

In this work, we use a combination of available genome sequences, manual curation, genome and transcriptome data, phylogenetics, expression profiling, and gene knockdown to investigate a family of genes from the CD36 superfamily in insects. The CD36 superfamily codes for membrane-bound scavenger proteins with a variety of ligands and signalling functions. In insects, the best characterized of these functions relate to signalling in response to environmental cues, such as diet, stress, innate immunization and chemoreception. There are a total of three CD36 families and here we focus on the sensory neuron membrane proteins (SNMPs) ^1^. The first SNMPs were identified from Lepidoptera ^2–4^ with SNMP1 expression shown to be highly expressed in pheromone receptor neurons of trichoid sensilla ^4^. Further evidence indicates that SNMP1 in *Drosophila melanogaster* is a co-factor along with DmelOR67d/Orco of the sex pheromone detection system and, therefore, critical for species-recognition ^5^. In addition, it was recently shown that SNMP1 interacts directly with DmelOR22a, a cis-vaccinyl acetate receptor, rather than Orco using Förster resonance energy transfer (FRET) system ^6^. The SNMP2 protein has yet to be fully characterized, but moth SNMP2 is known to be expressed in supporting cells around odorant sensitive neurons (OSNs) in olfactory antennae sensilla ^2,7^. Recent studies also showed that the moth SNMP2 is broadly and highly expressed in antennae, legs and wings ^8–10^. However, *Drosophila* transcriptome profiling from FlyBase has shown SNMP2 has limited expression in the gut.

In our present work, the objective was to characterise the SNMP family in moths, but an advantage is that we began with phylogenomic approach in order to determine the true orthology of SNMP genes. Therefore, we first fully ascertained the SNMP family and generated a robust and accurate phylogenomic model. As a result, we discovered that a novel SNMP gene arose in Lepidoptera and named SNMP3. We then proceeded to integrate public transcriptomic evidence to build hypotheses around function. Second, by integrating the expression data with the classic phylogenetic clustering, we showed that the family has been misannotated, likely due to previous dependence on BLAST similarity searches rather than a phylogenomic approach. Third, we provided multiple lines of evidence including RNAi, split-ubiquitin yeast hybridization system, and expression data, to show that - in spite of its sequence divergence - the function of the SNMP1 complex is conserved between Diptera and Lepidoptera, and is essential to mating recognition. We recommend it as a model for studying protein co-evolution driven by assortative mating.

## Results

### Full ascertainment of SNMPs in various insect species

Using official genome annotations, community contributions and similarity searches we collected a total of 81 SNMP genes from 35 insect species as distributed amongst seven orders (Table 1). In order to fully ascertain the gene family, we manually curated each species for which we had transcriptome or genome assemblies available (Supplementary Table 1), creating a high confidence dataset of 19 species from all seven orders. We also searched two non-insect invertebrate species (*Daphnia pulex* and *Caenorhabditis elegans*) but we did not find any SNMP genes that could be unambiguously inferred as orthologous to our insect core set. Of these 81 SNMP genes, 53 had accurate public gene structures; we edited four structures, contributed a further 20 new annotations and report four partial genes found in public transcriptome databases (Table 1). Overall, 19 of the species had full ascertainment for the SNMP gene family (i.e. the correct public annotations for all SNMPs orthologues known for each order) and 16 had some missing or incorrect annotations (Table 1). It is notable that in spite of available full genome sequence, we did not identify the homologue of *Apis mellifera* SNMP2 in *Nasonia vitripennis* and *Solenopsis invicta*. In addition, *N. vitripennis* had four tandemly duplicated *SNMP1* genes (XP_001606602, XP_001606682, XP_001606675, XP_001606692), which we could not verify due to the lack of primary data for the assembly. Further, the pea aphid (*Acyrthosisphon pisum*) had only one SNMP gene family member, even though another Hemipteron with a full genome sequence (bedbug, *Cimex lectularius*) had three gene family members. Finally, we annotated a new insect order, Anoplura, using the published genome sequence of body louse (*Pediculus humanus corporis*) ^11^, identifying two SNMP genes. Due to the lack of RNA-seq or comparative data, however, we were not able to characterize the full length of these genes.

**Table 1.**
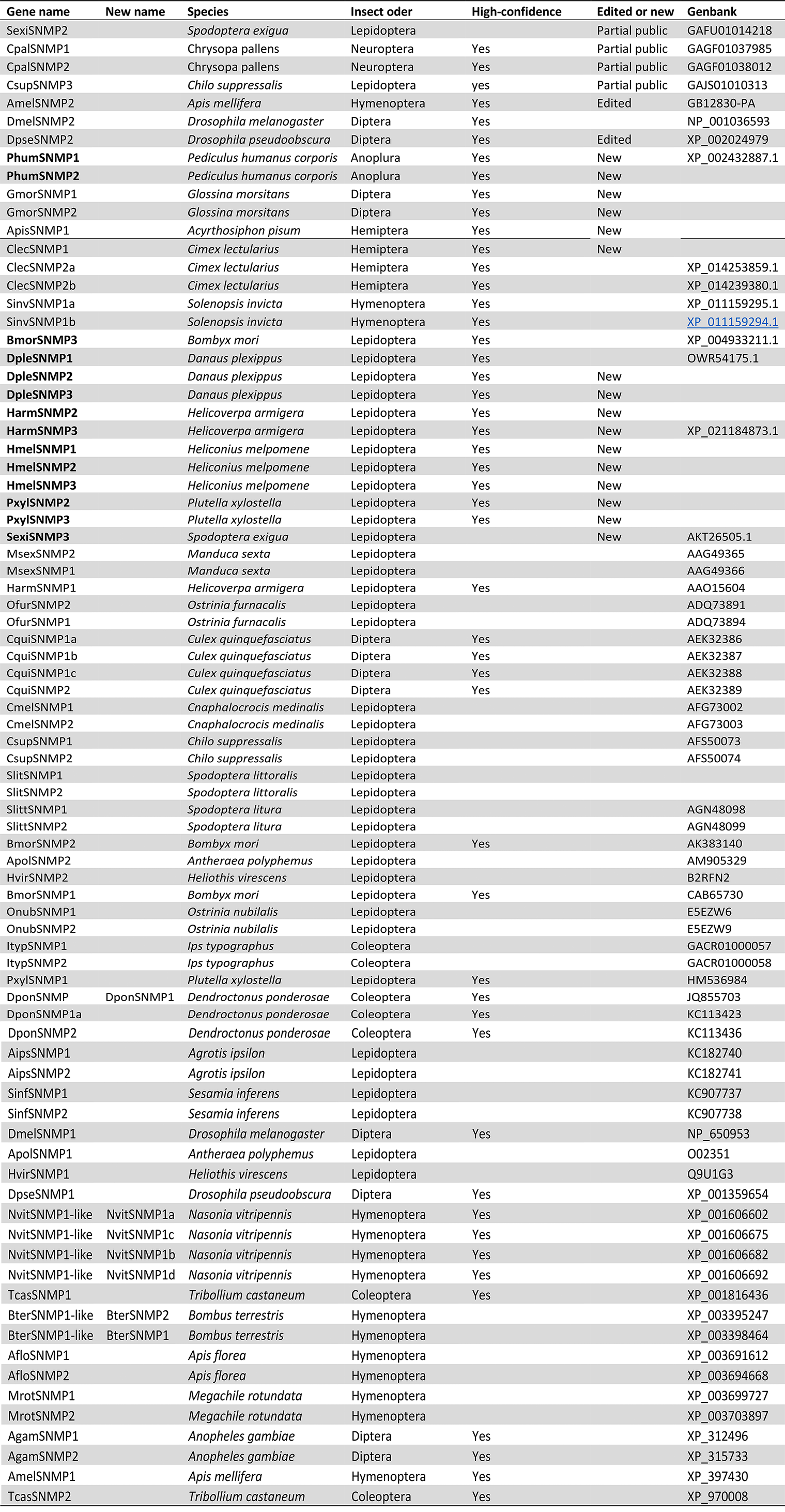
Identification of insect SNMPs.

Guided by available evidence, we edited two genes. Using the *D. melanogaster* homologue from FlyBase (FBgn0035815) we identified the *D. pseudoobscura* orthologue via OrthoDB (cluster EOG79D9HT) and named *DpseSNMP2* (FBgn0080333; XP_002024979), which had two extraneous, unsupported exons (4^th^ and 5^th^). For *Apis mellifera* (Hymenoptera), we used tBLASTn with *DmelSNMP2* and found that the *A. mellifera* RefSeq annotation of *SNMP2* (XP_001121085) was incorrect; we built a new gene based on GB12830-PA by identifying the missing N-terminus and a middle exon. For *Solenopsis invicta* (Hymenoptera) we used the genome raw data (GCA_000188075.1) to annotate and manually extend two hypothetical proteins (EFZ18285 and EFZ18363), which are tandem arrayed in scaffold_02797 (Table 1).

Further, we identified a number of new genes in Anoptera, Lepidoptera, Diptera and Hemiptera: For *Pediculus humanus corporis* (Anoptera) we found two SNMP genes identified from the publicly genome sequence available from Ensembl (scaffolds DS235882 and DS235862). Without access to RNA-Seq data, both gene annotations are missing their C-termini and SNMP1 is missing the N-terminus.

For *Cimex lectularius* (Hemiptera) we found a total of three SNMPs using a genome assembly produced by the Baylor College of Medicine and released under the Fort Lauderdale agreement (GCA_001460545.1; https://apollo.nal.usda.gov/cimlec/sequences). The genome had significant misassemblies. By aligning the raw assembly DNA-seq data to the genome assembly we were able to identify whether regions of concern may be linked to assembly issues. The full-length sequence from SNMP1 (512 amino acid) was annotated in scaffold 41 and verified with an RNA-seq assembly. Since our work, a new assembly (GCA_000648675.3) was produced with a more correct gene model (XP_014251146) however our 5’ end of gene. Our phylogenomic approach was not conclusive which gene model was more correct. but For SNMP2a, a partial gene encoding 448 amino acids was identified from scaffold_56 and the DNA- and RNA-Seq data point towards a misassembly for the last exon. Finally, a full-length SNMP2b encoding 510 amino acid was curated from scaffold_1 and verified with an RNA-seq assembly. For *Acyrthosiphon pisum* (Hemiptera): we searched the second genome version (GCA_000142985.2). We identified the complete CDS of SNMP1 from scaffold EQ121771 but did not identify any sequences homologous to the other SNMPs.

For *Glossina morsitans* (Diptera) we used the tsetse fly genome (https://pre.vectorbase.org/) that is currently available from VectorBase and identified a partial SNMP1 and a full ORF of SNMP2 orthologues from scaffolds scf7180000644980 and scf7180000648879 respectively. Transcriptome data was available via VectorBase’s Sanger EST transcripts but we found support only for the last exon of SNMP2 (FM961280.1).

For *Bombyx mori* (Lepidoptera) we found a novel gene in scaffold DF090341.1, which was named *SNMP3* here. Two automated predictions in SilkDB were in that region (BGIBMGA012262 and BGIBMGA012263) but classified as ‘antigens’. KAIKObase had three automated annotations (KAIKOGA048486, KAIKOGA048483 and KAIKOGA048479) but none was accurately reflecting the correct gene model. We annotated the full gene and verified it was a real SNMP homologous to the CD36 family, aligning with the *B. mor*i and other Lepidopteran SNMPs. We verified the structure using an EST sequence from KAIKObase (FS809883) spanning the exons of the two automated annotations; the combined size (508 amino acids) is similar to what is expected of SNMPs. We tested our results using primers designed from the annotated sequence and amplified a RT-PCR product of the expected length.

For *Helicoverpa armiger*a (Lepidoptera), we used a recently published whole genome ^12^ and discovered a new SNMP gene which was orthologous to the *BmorSNMP*3. The automated predictions had three genes spanning that area, but Sanger EST assemblies from InsectaCentral (IC29058AcEcon7899, IC29058AcEcon17308, IC29058AcEcon21585) ^13^ and RNA-Seq spanning exon-intron junctions allowed us to annotate the gene model. Further, we verified the gene models of previously reported *SNMP1* (AAO15604) and *SNMP*2 (Supplementary Table 2). For all three genes, we verified the expected length and expression using RT-PCR.

For *Spodoptera exigua* (Lepidoptera): the *SNMP3* homologue was identified by searching GenBank and reassembling previously reported cDNA sequences (GAFU01000186 and GAFU01009183). Further, we identified a partial *SNMP2* gene from GAFU01014218.

For *Plutella xylostella* (Lepidoptera) we found homologues for *SNMP2* and *SNMP3* by manual curation of the previously published *P. xylostella* genome (GCA_000330985.1). A 523 amino acid long *SNMP2* was located on scaffold KB207336.1 and was curated with the assistance of transcriptome data (accessions PXUG_V1_014487, PXUG_V1_12791 and RS_pxwb_0023634_c0_s001) available from the genome project databases (http://iae.fafu.edu.cn/DBM; http://dbm.dna.affrc.go.jp/px). The 541 amino acid homologue of *SNMP3* was located on scaffold KB207313.1 and also supported by RNA-Seq evidence, which was however partial due to assembly gaps or a misassembly. We used profile-guided alignments with other lepidopteran *SNMP3* genes and the assembled transcriptome (PXUG_V1_028331) to extend the gene to its full length. We further verified the accuracy of this sequence by performing a Trinity denovo assembly of all the raw RNA-seq data: a 2451 bp contig verified the start codon, stop codon, 5’ and 3’ UTR sequences.

For *Danaus plexippus* (Lepidoptera) we found a total of three full-length SNMPs using genome sequence (GCA_000235995.1). The *SNMP1*, *SNMP2* and *SNMP3* genes were initially annotated from scaffold JH380873, JH384389 and JH390655, respectively. As the scaffold JH384389 was very short (5 kb), we re-annotated using a third version of the *D. plexippus* assembly (scaffolds DPSCF300414, DPSCF300039 and DPSCF300068 respectively) which was available only from MonarchBase directly (http://monarchbase.umassmed.edu). Final full-length sequences were curated, encoding 519 AA, 524 AA and 522 AA, respectively. We used the publicly available RNA-Seq data (SRR585568) to verify all but one of the intron/exon junctions. The only notable discrepancy was for the *SNMP1* gene for which we do not have RNA-Seq support for the 6^th^ exon, yet the RNA-Seq do link exons 1-5 and 7-9 and a multiple sequence alignment versus of all lepidopteran *SNMP1* supports the 6^th^ exon. Further, the RNA-Seq data provided evidence that the underlying genome sequence had a frameshift-causing insertion at position 768,105.

For *Heliconius melpomene* (Lepidoptera), we identified and annotated all three SNMPs using the published genome sequence (GCA_000313835.2). The SNMP1 gene was located on scaffold HE671626 and an automated prediction had partially annotated the gene. Using the *Danaus plexippus SNMP1* gene we conducted tBLASTx searches to identify one N-terminus exon but due to gaps, putative misassemblies and lack of RNASeq from relevant tissues, we failed to identify the full C-terminus for this species. The full SNMP2 and SNMP3 genes were similarly curated on scaffolds HE671451 and HE672039 respectively. In the case of *SNMP3*, the gene model was supported by RNASeq evidence.

#### Phylogenetic characterization

In the phylogenetic analysis, the insect SNMPs form an independent group from CD36 genes (Figure 1A). Furthermore, insect SNMPs shows that three distinct clusters are formed: the SNMP1 clade, SNMP2 clade (comprising of the Hymenoptera and Lepidoptera SNMP2) and SNMP3 clade (containing the *Drosophila* SNMP2). Within each of these clades, each taxon group clusters separately, indicative of an evolutionary process that is broadly stable but there are notable exceptions.

**Figure 1.**
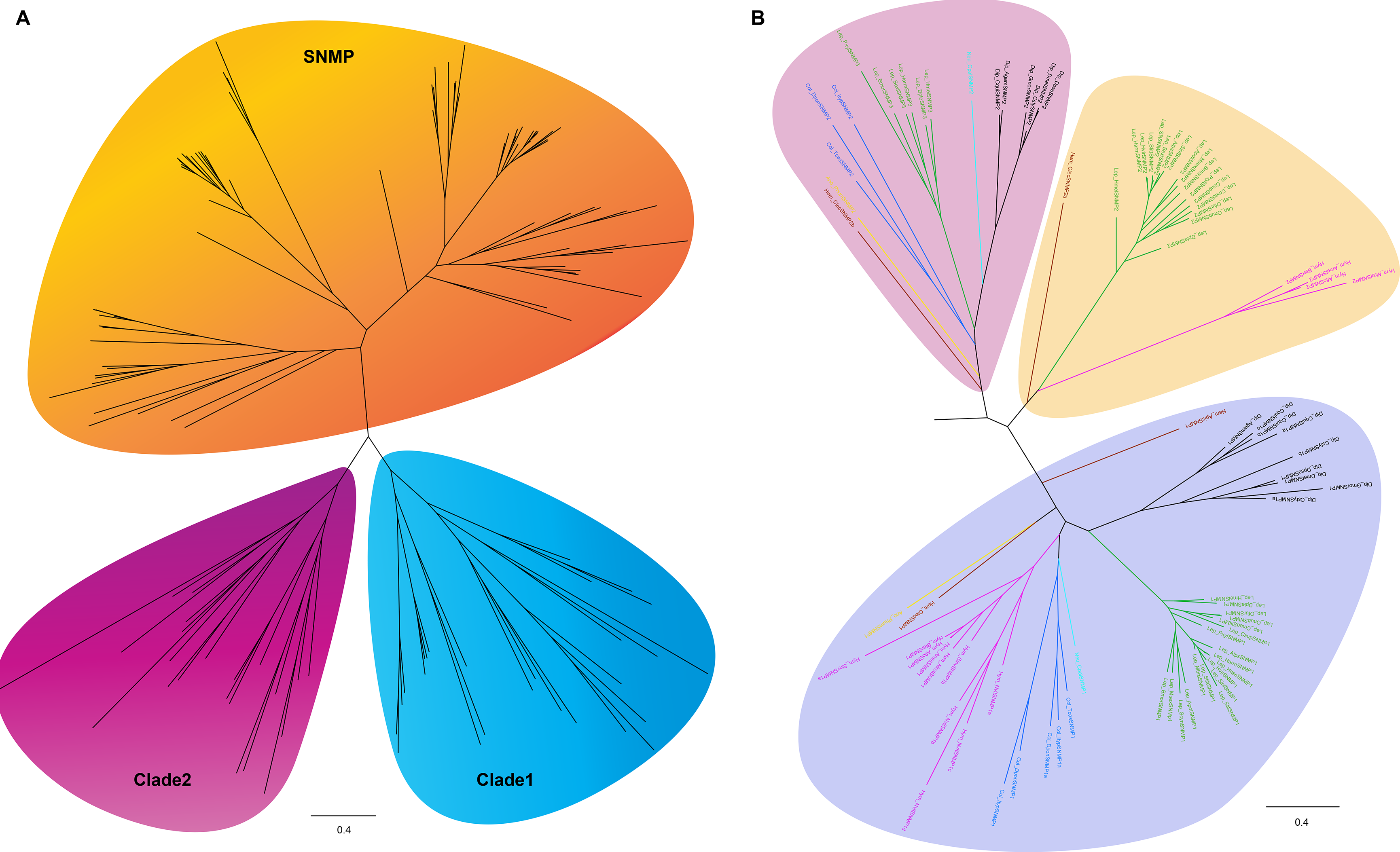
Phylogenectic analysis of insect SNMP/CD36 homologs. A, Phylogenetic analysis of insect SNMPs and CD36 homologs. Blue, green and red shade present clade 1, 2 and SNMPs. B, SNMPs fall into three sub-clades (SNMP1, SNMP2 and SNMP3), which have been highlighted in yellow, green and blue. B, Neigbhor-joining tree of insect SNMPs. Dipteran (black), Lepidopteran (green), Hymenopteran (Fuchsia), Hemipteran (Violet), Anopteran (Olive) and Neupteran (Aqua) and Coleopteran (Blue) SNMPs were labelled. To simplify, abbreviation name for each insect species are listed as below: *Helicoverpa armigera*, Harm; *H. assulta*, Hass; *Heliothis virescens*, Hvir; *Mamestra brassiae*, Mbra; *Antheraea polyphemus*, Apol; *B. mori*, Bmor; *Samia cynthia ricini*, Scyn; *Manduca sexta*, Msex; *Chilo suppressali*s, Csup;*Cnaphalocrocis medinalis*, Cmed; *Ostrinia nubilalis*, Onub; *Ostrinia furnacalis*, Ofur; *Plutella xylostella*, Pxyl; *Agrotis ipsilon*, Aips; *Danaus plexippus*, Dple; *Spodoptera litura*, Slit; *S. frugiperda*, Sfru; *D. melanogaster*, Dmel; *D. pseudoobscura*, Dpse; *A. gambiae*, Agam; *A. mellifera*, Amel and *T. castaneum*, Tcas. The SNMP/CD36 in *D. melanogaster*, *D. pseudoobscura*, *A. gambiae*, *A. mellifera* and *T. castaneum* are identical to those used previously ^22^ except few renamed because of new released genome/transcriptome version. The ML tree was built by RAxML. The bootstrap is 1000 replications.

An obvious feature in this phylogeny is that what has been called a Lepidopteran SNMP2 is not actually orthologous to the *Drosophila* SNMP2. When SNMP2 was discovered, researchers seem to have named it using the best BLAST score against *Drosophila* SNMP2 rather a reciprocal BLAST approach or the extensive phylogenetic approach we have done here. The lack of functional studies is common in similar publications but with the innovations in genome sequences or RNAi methodology, we feel it is easier than ever to provide further evidence. The new gene we discover here, named by convention as SNMP3, is more likely to be the true orthologue of the *Drosophila* SNMP2. This SNMP3 exists in all Lepidoptera species we surveyed and is monophyletic.

#### Comparison of time- and spatial-specific expression of SNMP orthologues in insects

The first SNMP was characterized from the antennae of wild silkmoth *Antheraea polyphemus* in 1997 ^4^. Subsequently, another SNMP homolog, named SNMP2, and more orthologs were identified from more Lepidoptera, Diptera, Coleoptera and Hymenoptera ^1^. To investigate whether SNMP orthologs share similar functional characteristics during development, semi-qRT-PCR was primarily performed to analyse the expression of SNMPs in multiple tissues of *B. mori*, *H. armigera* and *D. melanogaster*, typical insect models of Lepidoptera and Diptera.

By comparing the expression of SNMPs from a number of tissues from both *B. mori* larvae and adults, we found that *BmorSNMP1* and *BmorSNMP3* have a more restricted time- and spatial-expression pattern (Figure 2A). As shown in Figure 2A, *BmorSNMP1* had expression limited in the adult moth antennae. Meanwhile, the expression of *BmorSNMP3* was strongly detected in larvae and adults midgut with lower expression in male head and slight expression in female head. Conversely, there is a wild range of expression of *BmorSNMP2* in all tested tissues during development.

**Figure 2.**
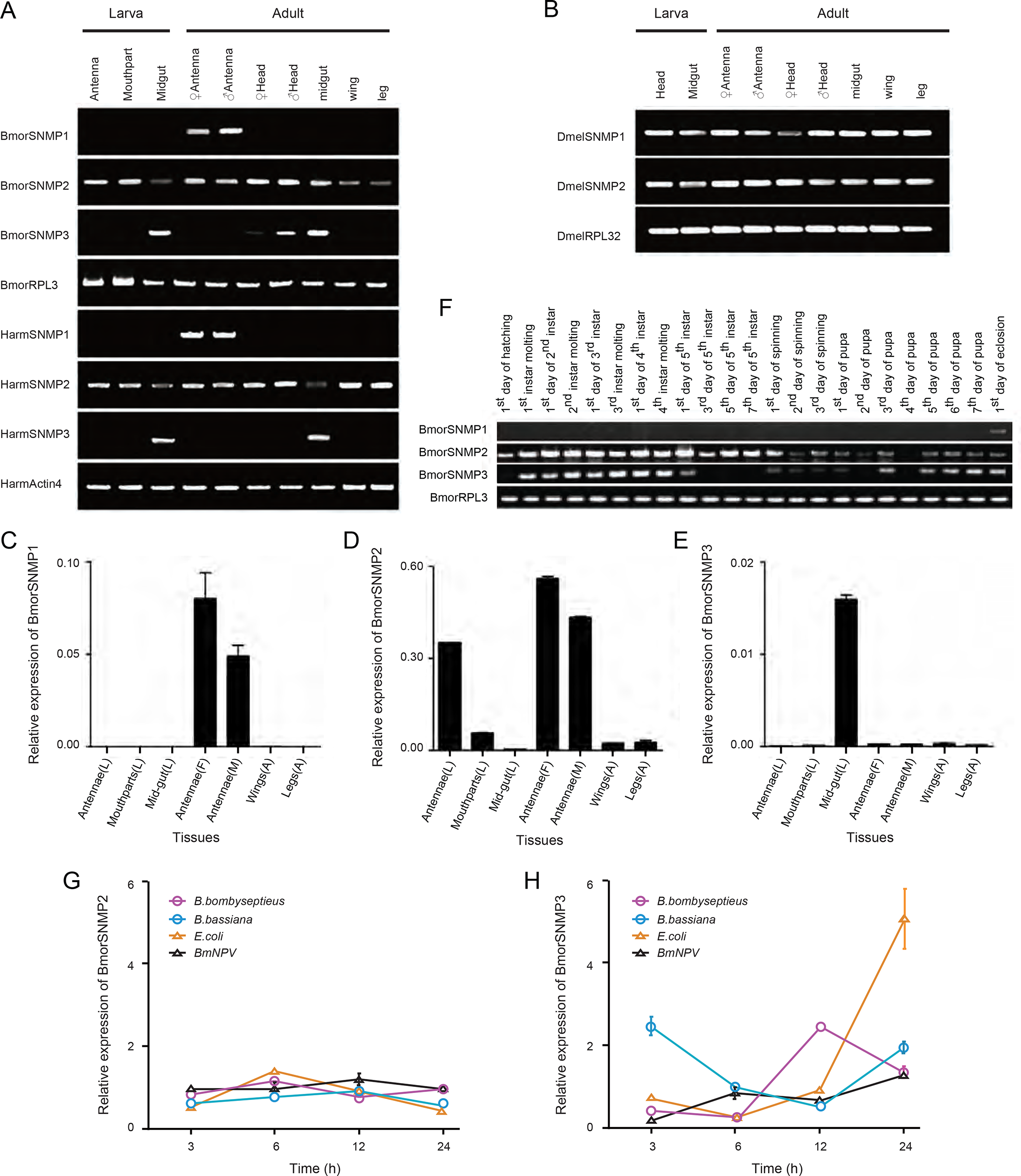
Specific tissue and time expression pattern of SNMPs. A, Gene expression of *B. mori* and *H. armigera* SNMP orthologs in multiple tissues by using RT-PCR. B, Expression of DmelSNMPs in *Drosophila* larval and adult tissues. C, D and E, the relative expression of BmorSNMP1, 2 and 3 in various larvae and adult tissues by using qRT-PCR. F, The expression of BmorSNMP1, 2 and 3 in different development stages from egg hatching to adult eclosion. G and H, i*n silico* expression ratio analysis of BmorSNMP2 and 3 in individual larva infested by bacteria or virus. The larvae at 3^rd^ day of 5^th^ instar were infected by *Bacillus bombyseptieus* (blue line with hard circles), *Beauveria bassiana* (purple line with hollow circles), *Escherichia coli* (black line with hard triangles) or *B. mori* nucleopolyhedrovirus (BmNPV) (orange line with hollow triangles) for three hours and then fed by normal artificial food. The whole insects were collected at different time for oligonucleotide arrays. The expression ratio of BmorSNMP2 and 3 were screened through microarray data using gene ID in genome.

In comparison with the expression in *B. mori*, similar patterns across tissues and development was found in *H. armigera* SNMPs (Figure 2A). Strikingly, *HarmSNMP3* gene was specifically expressed in both larvae and adults midgut. Using RNA-seq data from the *H. armigera* Genome Consortium, we found that *HarmSNMP1* was also expressed in male adult tarsus and 5^th^ instar antennae but our semi-qRT-PCR did not capture that. *HarmSNMP2* was also expressed in abdomen, thorax, tarsus, testes, prepupae, and larval foregut, antennae, mouthpart, cuticle and salivary gland besides adult antennae and head (with antennae) according to RNA-seq data which is supported from our own PCR result showing broad expression. For *HarmSNMP3*, the RNA-Seq data showed that it was expressed in the midgut, as supported by our RTPCR, but also embryos.

Compared with *B. mori* and *H. armigera*, the expression profiles of the *SNMP1* gene in *Drosophila* is distinct. *DmelSNMP1* and *DmelSNMP2* shared the similar expression characteristic of high expression in multiple tissues of larvae and adults (Figure 2B). Those are supported by previous work on DmelSNMP1 (c.f. FlyBase FBgn0260004 and relevant modENCODE links) that show that it is highly expressed in adults including carcass, eye, fat body, head, hindgut, midgut, salivary gland and larval trachea. *DmelSNMP2* (c.f. FBgn0035815 and relevant modENCODE data) is widely expressed across development but with higher expression in crop and hindgut compared to adult carcass, head, larvae hindgut, and salivary gland.

We further performed qRT-PCR on BmorSNMPs in larval antennae, larval mouthparts, larval midguts, female adult antennae, male adult antennae, adult wings and adult legs. The results (Figure 2C, D and E) are consisted with the RT-PCR results (Figure 2A and B). We also studied the time-specific expression of BmorSNMPs from the egg hatching to the adult eclosion (Figure 2F). Our results showed that BmorSNMP1 is only expressed when adult eclosed from pupae stage (Figure 2F). However, BmorSNMP2 and BmorSNMP3 are expressed in all stages from egg hatching to adult eclosion. Interestingly, BmorSNMP2 and BmorSNMP3 showed higher expression at larval stage than pupae stage, suggesting they may play important roles at larvae stage.

##### SNMPs play a role in immunity responses

To explore whether BmorSNMPs are involved in immune response, public microarray data (http://www.silkdb.org/silkdb/) were used to check the expression ratio of genes after the larvae were infected by *Bacillus bombyseptieu*s, *Beauveria bassian*a, *Escherichia coli* or *Bombyx mori* nucleopolyhedrovirus (BmNPV). When compared to a control, the ratio of expression for *BmorSNMP2* varied only slightly between types of inoculations and time after inoculation (Figure 2G), an indication that SNMP2 is not involved in immune response. For *BmorSNMP*3, however, the ratio of expression was notably increased during the first three hours after *B. bombyseptieu*s inoculation and then decreased (Figure 2H). After inoculation with *B. bassian*a, the ratio peaked the 12th hour. After inoculation by BmNPV, the expression ratio was elevated 12 hours post induction and reached its maximum at 24h, which was the total length of the experiment. For *E. coli* there was no variation (Figure 2H). These expression rate changes as a response to infection and that difference across types of infections indicates that SNMP3 is involved to immune responses. Due to lack of any available probe, we could not analyze the expression ratio changes of BmorSNMP1 but we believe it is unlikely that a gene that is specifically expressed in adult antennae is involved in immune response.

### Functional characterization of the silkworm SNMP1 involved in sex pheromone perception

To compare the function of BmorSNMP1 in pheromone detection, pheromone receptor BmorOR1 was also investigated in this study. BmSNMP1 and BmOR1 showed similar expression profile in male antennae from 4^th^ day of pupa using RT-PCR (Figure 3A). It provides cue for the time of dsRNA injection.

**Figure 3.**
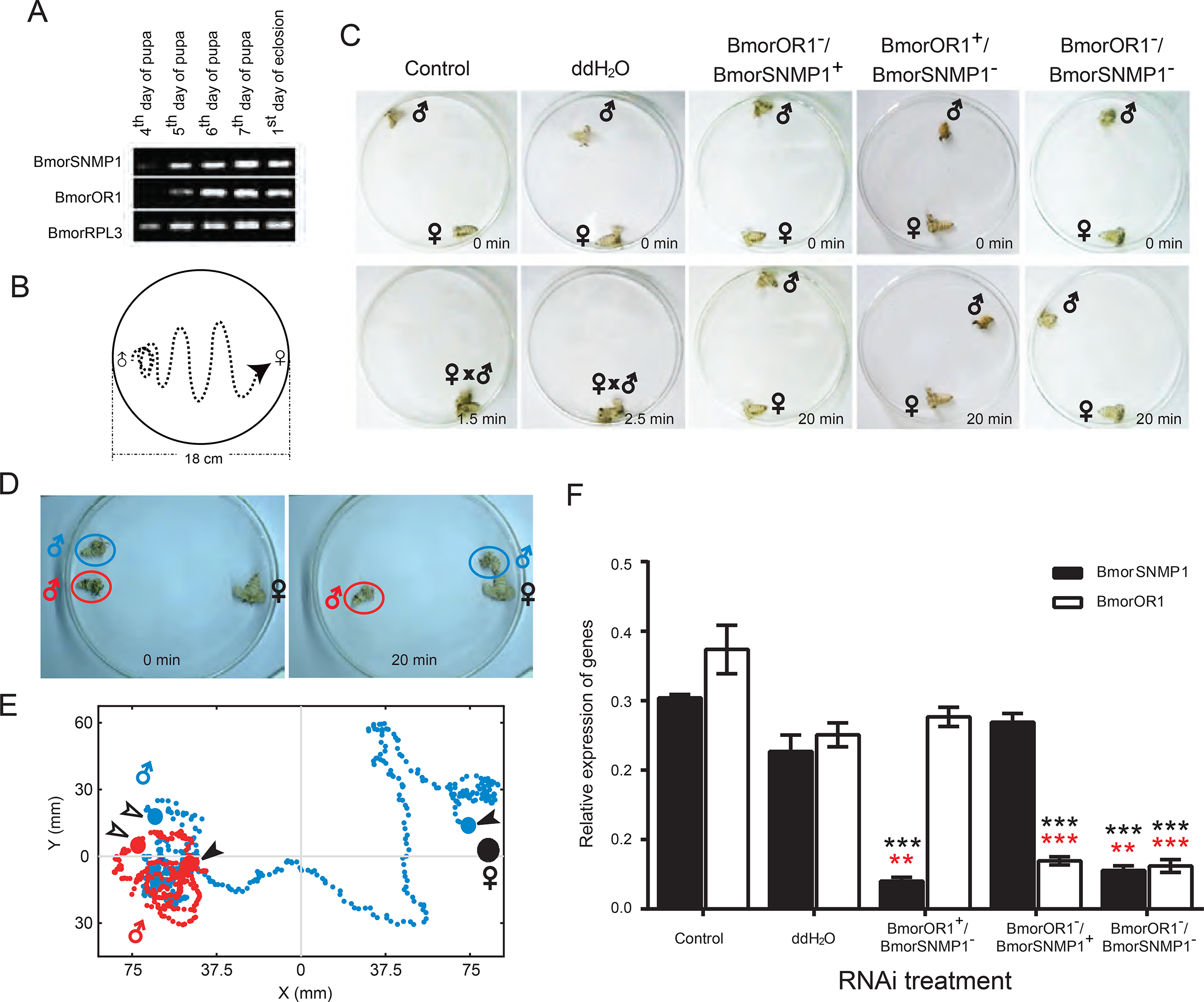
Behaviour assays on SNMP1/OR1 knock-down moth treated by RNAi. A, Expression of BmorSNMP1 and BmorOR1 in male antennae during the last four days of pupae and the 1^st^ day of the eclosion. B, Schematic diagram of one male-female moth recognizing and finding assays. C, Positions for single pair of male-female moth before and after 20 min. D and E, Photo and trace examples for competing behavioural assay. Red and blue round bars present intact (control) and RNAi treated male moth respectively. Red and blue dot lines indicate moving paths of moth getting to female in 3 min. Hollow and hard arrows respectively suggest start-stop positions. F, Relative expression level of BmorSNMP1 and BmorOR1 in male moths’ antennae after RNAi were detected by real-time PCR. Hard black and hollow volume indicated the expression of BmorSNMP1 and BmorOR1 respectively. Error bar showed the standard error of the mean, SEM. Data were also analyzed by using two-tailed Student’s t-test. Black stars presented the difference between RNAi treatment and control, while red stars indicated that between RNAi treatment and water. \**p*< 0.05, \*\**p*< 0.01, \*\*\**p*< 0.005.

Behavioural assays were performed within 48 hours after eclosion. In one male-female test group, the position of moths was shown (Figure 3B) with the most amount of possible space between them in the dish. Generally, behavioural response could be elicited by female in the majority of male moths. This behaviour ranges from wing vibration and anemotactic walking to female tracking the source of pheromone (Supplementary Video 1). The time for male from recognizing to finding females is postponed after RNAi knocked down of the expression of either BmorSNMP1 or BmorOR1 or both (Figure 3C, Supplementary Video 2). In control, male moths fluttered wings “madly” and showed anemotactic towards the female (Supplementary Video 1). All males can recognize and reach the female within 10 min with average time 160 sec. 93% of ddH2O treated males could find female by slightly delayed with 250 sec of average time. 87% of male could find female in the first 10 min. 6 % could locate female in the next 5 min. However, 2 % of males couldn’t find the female. On the contrary, RNAi treatment results in ≤ 30 % males finding female in the first 10 min and ≥ 34 % of males could not locate the females within the 20 min though some behavioral responses such as fluttering wings could be elicited by female (Supplementary Video 3). The average times of those could locate female after RNAi treatments are around 524 sec (BmorSNMP1^−^), 620 sec (BmorOR1^−^) and 641 sec (BmorSNMP1^−^/BmorOR1^−^) respectively.

For further comparison of RNAi treated and control males, competing behavioral assays were performed by placing a pair of male moths in the same arena with one female at the longest diameter (Figure 3D and E). Both males exhibit fluttering wings which is a typical sexual behavioral indicating both males detected the pheromone elicited by the female (Supplementary Video 4). However, only control males successfully moved towards and located females within 3 min though the control males did spend almost the 1 min pursuing the RNAi male. A similar result was found for BmorOR1 RNAi knocked down males with only control moths locating the female within the first 3 mins (Supplementary Video 5). The rate for non-eclosion pupae is 9-10 % after dsRNA injection comparing with 5 % for ddH_2_O injection and 0 for control.

After behavioral testing, antennae were collected from the same moths and the expression level of BmorSNMP1 and BmorOR1 in the antennae was detected by real-time PCR. The expression level of each gene has been statistical analyzed by Two tailed and Unpaired Student t-test. The results revealed that expression of both BmorSNMP1 and BmorOR1 were not significantly different between control and ddH_2_O injection (Figure 3F). Further BmorSNMP1 dsRNA did not affect the expression of BmorOR1 and vice versa. However, a significant down-expression of BmorSNMP1 (t=46.56, df=4, *p*<0.001) and BmorOR1 (t=8.59, df=4, *p*<0.001) after RNAi treatment have been found comparing with control and reduced remarkably, BmorSNMP1 (t=8.14, df=4, *p*<0.01) and BmorOR1 (t=9.96, df=4, *p*<0.001), comparing with ddH_2_O injection. In binary RNAi treatment, both BmorSNMP1 (t=29.89, df=4, *p*<0.001) and BmorOR1 (t=8.62, df=4, *p*<0.001) were significantly decreased compared to control as well as ddH2O injection (tSNMP1=6.90, df=4, *p*<0.01; tOR1=9.64, df=4, *p*<0.001).

#### SNMP1 interacted with olfactory receptors

Previous studies mentioned DmelSNMPs would be involved in olfaction and DmelSNMP1 has a potential interaction with DmelOR22a, the pheromone receptor forcis-vaccinyl acetate ^6^. We postulated that there is a crosstalk between insect SNMP1s and pheromone receptors (PR) or Orco homologs in promoting pheromone response. To investigate this, split-ubiquitin yeast hybridization was performed which is considered as a promising method to reveal the interaction between transmembrane proteins. It was demonstrated that both SNMPs and ORs have cytosolic N-terminus which provide cues for choosing vectors for yeast hybridization. pPR3N vector was chosen for SNMP1s and BmorOR2 as the prey constructs versus pBT3STE vector for pheromone receptors as bait constructs. The interaction between pNubG-Fe65 and pTSU2-APP was used as positive control (Figure 4A). To exclude the internal interaction between vectors, the interaction of empty constructs of pPR3N and pBT3STE was shown and treated as negative control.

**Figure 4.**
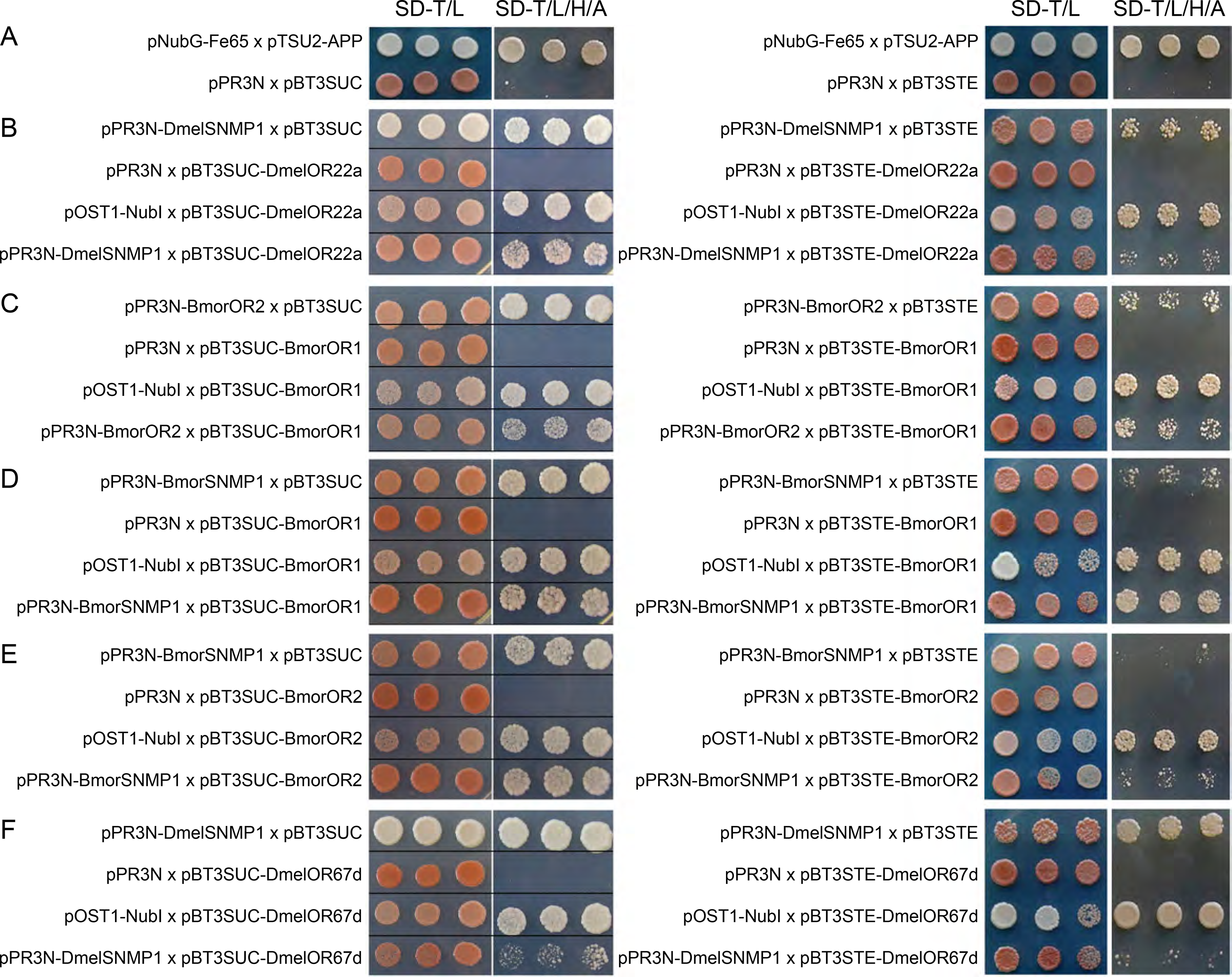
The interaction between SNMP1 and pheromone receptor from *D. melanogaster*, *B. mori* and *H. armigera* using split-ubiquitin yeast hybridization system. After co-transformed two fusion protein constructs into yeast, positive clones could be shown in amino acid defective selection medium (SD-T/L or SD-T/L/H/A) if the two target proteins interacted with each other. A, two pairs of pNubG-Fe65 x pTSU2-APP and pPR3N x pBT3STE were used as positive and negative controls respectively. B, the reported interaction of DmelSNMP1 and DmelOR22a was examined by yeast hybridization assays. C, the interactions between the BmorOR2 (Orco) and pheromone receptor (BmorOR1). D and E, the potential interactions of SNMP1 orthologs with pheromone receptors and Orco in *B. mori* were separately evaluated by identical treatment. F, the interactions between the DmelSNMP1 and DmelOR67d.

Due to the clear interaction of DmelSNMP1 (CG7000) with DmelOR22a (NM_078729) has been argued, this pair was tested primarily which would provide evidence to whether the yeast system is suitable to our investigation. After 4-day incubation of yeast co-transformed pPR3N-DmelSNMP1 with pBT3STE-DmelOR22a, the positive results have been shown by both SD-Trp-Leu (SD-T/L) and SD-Trp-Leu-His-Ade (SD-T/L/H/A) defective selection (Figure 4B). These results confirmed that DmelSNMP1 interacts with DmelOR22a which has been reported ^6^, indicating the split-ubiquitin yeast hybridizaiton system could approach our purpose. Subsequently, to evaluate the interaction between SNMP1 othologs and pheromone receptors in each insect species, the binary pairs of DmelSNMP1/DmelOR67d and BmorSNMP1/BmorOR1 have been detected in yeast hybridization system. The results were suggestive of the interaction of BmorSNMP1/BmorOR1 (Figure 4D) and DmelSNMP1/DmelOR67d (Figure 4F). Moreover, to elucidate whether BmorOR1 and BmorSNMP1 could separately interact with Orco homolog, we further identified co-transformed BmorOR2 (Orco) with BmorOR1 or BmorSNMP1 into yeast. Expectedly, positive results have been revealed (Figure 4C and E). Finally, β-galactosidase activity was evaluated for examining the intensity of the pairwise interaction (Supplementary Table 5). Statistical analysis indicated that there was slight higher but no significant difference in each pair compared with negative control. Taken together, those results inferred that SNMP1 orthologs interact with pheromone receptor/Orco. However, the pairwise interactions are rather weak which did initiate the expressions of HIS3 and ADE2 report genes but couldn’t trigger β-galactosidase activity in yeast.

To exclude the unfavorable impact of vectors on fusion protein expression, identical experiments as described as above have been practiced using pPR3N vector for pheromone receptors as the prey constructs versus pBT3STE vector for SNMP1s and BmorOR2 as bait constructs. Consequently, similar results could be found (data not shown). Finally, we proposed a new mode of moth Orco, SNMP1 and pheromone receptor (OrSo) function mechanism (Figure 5). Moth Orco, SNMP1 and pheromone receptor form a heteromer to function when the pheromone compound is delivered by pheromone binding protein in the sensilla (Figure 5).

**Figure 5.**
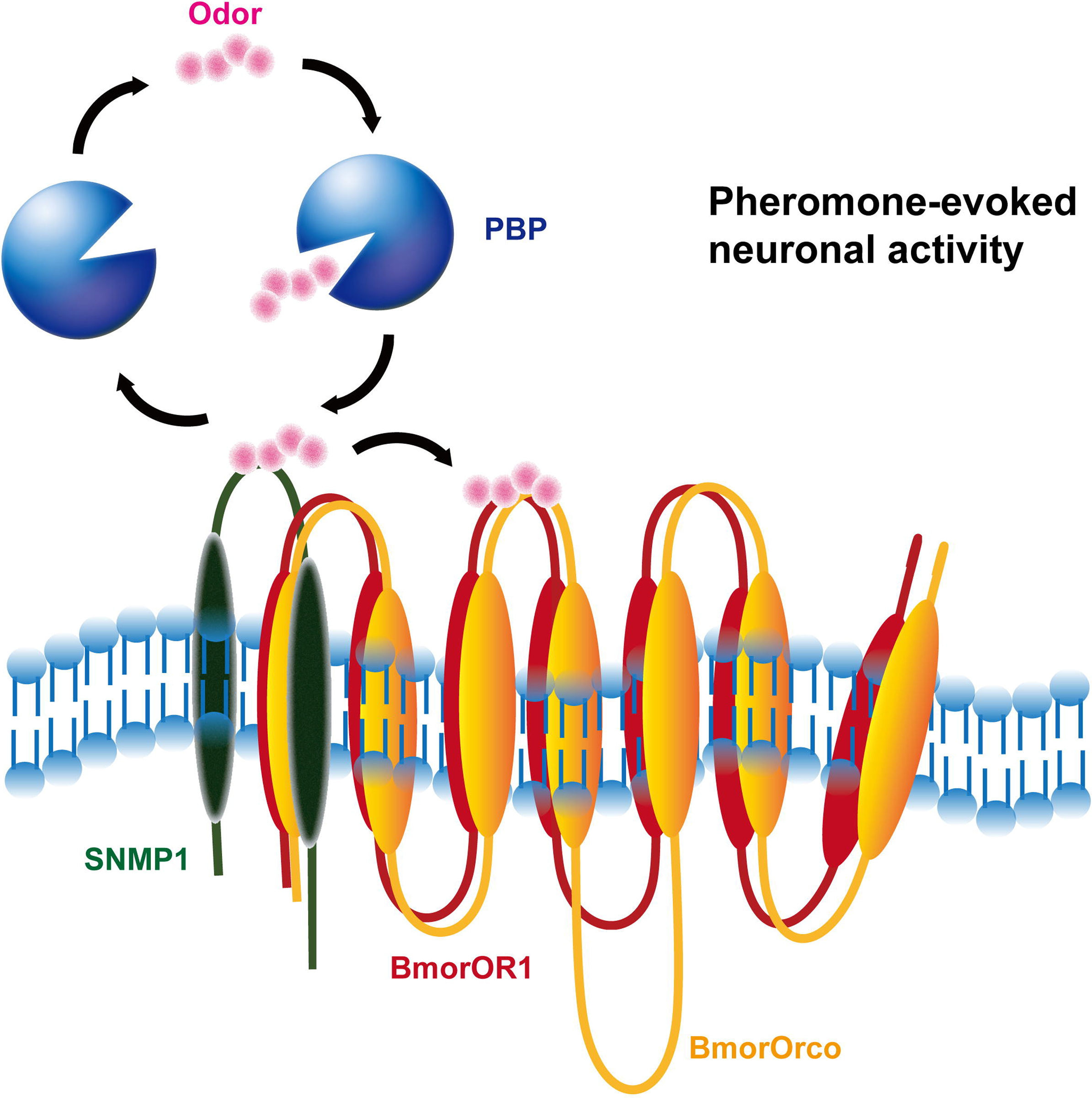
The new mode of Orco, SNMP1 and odorant receptor (OrSo) function mechanism.

## Discussion

Even though the SNMP subfamily has been known for years, it has not been well studied in Lepidoptera species. The first SNMP was characterized from the wild silkmoth *A. polyphemus* ^4^. Subsequently, draft whole genome sequencing (WGS) and shallow cDNA sequencing in *D. melanogaster* and other insects allowed the *in-silico* identification of a second protein, named SNMP2, from more Lepidoptera, Diptera, Coleoptera and Hymenoptera ^1–3,7^.

Due to the lack of functional work, the naming followed a practice standard in most whole genome sequencing (WGS) projects: first the genes are identified using BLAST searches and, assuming full ascertainment, a draft phylogenetic tree is constructed. Therefore these phylogenies are often derived on an average linkage clustering (based on sequence identity) or using an unrealistic evolutionary model. Rarely are the assumptions of the phylogeny reconstruction taken into account. Further, authors rely on bootstrap values as the sole index of accuracy and are rarely interested in investigating deeper. In the case of SNMPs, this network topology guided the assignment of orthology with gene naming following the convention of serially naming individual sequences as they were described in BLAST searches. Sadly this is still a standard practice in whole genome annotation but new automated procedures such as OrthoDB ^14^ have been made available to facilitate curators. These procedures still assumes full ascertainment and an accurate network topology are therefore best suited to be used in conjunction with curation. Further, these approaches neglect the use of expression profiling as a robust approach to assign function.

To date, SNMP1 expression was shown to be highly expressed in pheromone receptor neurons of trichoid sensilla ^3,7^. And SNMP1 may directly interact with odorant receptors rather than Orco in those neurons (Benton et al., 2007; German et al., 2012) suggesting SNMP1 may play a specific role in pheromone detection. Subsequent functional studies were limited to *Drosophila* and they showed that SNMP1 is essential for the pheromone receptor (OR67d) to detect the sex pheromone (11 *cis*-vaccenyl acetate) in a native environment ^5,15,16^. The second protein is yet to be thoroughly characterized that moth SNMP2 is expressed in support cells around odorant sensitive neurons (OSNs) in many olfactory sensilla of antennae ^2,7^. Recent study also showed that *Sesamia inferens* SNMP2 is broadly and highly expressed in antennae, legs and wings ^8^.

In this work, we undertook to fully ascertain the SNMP family in the lepidopteran model species the silkworm (*B.mori*) and all other insect with genomic or transcriptomic data. Using all available public data from insects we verified that a novel SNMP gene arose in Lepidoptera. Second, by integrating the expression data with the classic phylogenetic clustering, we show that the fruitfly SNMP2 clusters with the Lepidopteran SNMP3 and *Tribolium* SNMP2, not the Hymenopteran SNMP2. Rather Hymenopteran SNMP2 and Lepidopteran SNMP2 form a distinct monophyletic clade of a gene that is constitutively expressed. Third, based on current expression analysis, moth SNMP1 is restrictively expressed in adult antennae. SNMP2 is broadly expressed in multiple tissues. SNMP3 generally concentrates in larvae and adult midgut indicating the possible function in fatty acid transportation. Further, *B. mori* SNMP3 is also found in adult head (without antennae) and is differentially expressed in whole body when challenged with bacterial infections. On the contrary, the fruitfly SNMPs have similar expression pattern in all tested tissues. Finally, we showed that the silkworm SNMP1 is indeed a functional orthologue of the fruitfly equivalent using RNAi and bioassays. For the first time, we showed SNMP1 does not only bind to pheromone receptor (BmorOR1), but also the co-receptor, Orco. Based on this discovery, we brought out a new mode of insect pheromone receptor functional complex in silkworm, which is composed of BmorOrco, BmorSNMP1 and BmorOR1. Our results provide new evidence for SNMPs function *in vivo*. These can be readily applied to other insects that can mate in a laboratory setting.

## Material and Methods

### Insect materials

*B. mori*, Dazao strain, larvae and adults were reared on mulberry leaves at 25°C by the State Key Laboratory of Silkworm Genome Biology, Southwest University. Multiple tissues studied for RT-PCR and real-time PCR are listed as follows: antennae, mouthparts and midgut were dissected from larvae at 3^nd^ day in 5^th^ instar; Adult antennae, heads without antennae, midgut, legs and wings were dissected from emerging adults. For studying the expression of BmorSNMP1 and BmorOR1 male antennae were dissected from pupae or male moths from the 4^th^ day of pupa, the earliest day when we could get antennae, to the 1^st^ day of eclosion. All collected tissues were immediately immersed into liquid nitrogen and stored at −80 °C until use. *H. armigera* and *D. melanogaster* (Canton-S) were provided by CSIRO Ecosystem Sciences. Same tissues of *H. armigera* were collected at the similar age by using the same methods as those for *B. mori*. Heads and midgut of *D. melanogaster* were dissected from 3^rd^ instar larvae. The same tissues as those of moths were dissected from 4 days aged virgin flies.

### Gene identification and phylogenetic analysis

Reported sequences of SNMPs and CD36 family members from Lepidoptera, Diptera, Coleoptera and Hymenoptera were downloaded from NCBI or relevant sources ^1–4,7,17–22^. The sequences of SNMP/CD36 in *D. melanogaster*, *Anopheles gambiae*, *Apis mellifera* and *Tribolium castaneum* used here are identical to those used previously ^22^. We searched known insect genome and/or transcriptome (Table 1) using tBLASTn ^23^. Candidate genes were annotated manually in Geneious R7 created by Biomatters (http://www.geneious.com). Multiple transmembrane domains of receptors were predicted by TMHMM (http://www.cbs.dtu.dk/services/TMHMM-2.0/).

Amino acid sequences were aligned with MUSCLE ^24^ using default parameters. With R AxML v8.2.9 ^25^, the maximum likelihood (ML) tree was computed via rapid full analyses with the PROTGAMMAWAG substitution model and 1000 bootstrap replicates.

### Semi-qRT-PCR

Total RNA was extracted by trizol (Promega, USA) according to the protocol provided by the manufacturer. Total RNA was treated using DNase I, then quantified and qualified by NanoDrop ND-2000 (Thermo Scientific, USA). First strand cDNA was synthesized using SuperScript III First-Strand Synthesis Super Mix (Invitrogen, USA). Expressions of SNMPs in multiple tissues and during development were tested by semi-qRT-PCR. All primers for both semi-qRT-PCR are listed in supplementary Table 3.

### In-silico expression analysis

To gain insight into the function of SNMPs involving in immunization, microarray-annotated silkworm datasets (http://www.silkdb.org/silkdb) were screened with tBLASTn^23^. The regulated expression of genes in individual larva are denoted by ratio comparing the gene expressed level in intact larva.

### RNAi bioassay

RNAi was performed using dsRNA. The dsRNA of BmorSNMP1 is spanning the second transmembrane domain (309-520 AA) (Fwd 5’ GCTAATACGACTCACTATAGGGAGATACAATGGGATTAAGACGA 3’; Rev 5’ GCTAATACGACTCACTATAGGGAGATTTGGCTGGTTCTTGATT 3’), while that for BmorOR1 is spanning from the third to fifth transmembrane domains (109-255 AA) (Fwd 5’ GCTAATACGACTCACTATAGGGAGATCTTGTATTAACTGGTCGCTTCA 3’; Rev 5’ GCTAATACGACTCACTATAGGGAGATGGCTGGCTTTAGGTCTCG). All primers are fused by T7 promoter. dsRNA was prepared using RiboMAX Large Scale System-T7 kit (Promega, UAS) according to manufacturer’s instructions. 30 µg dsRNA in 10 µl ddH2O was micro-injected into one antenna on the 4^th^ day of pupae. Intact and ddH2O injected pupae are used as control. 50 male pupae were used for each test group. Chlortetracycin Hydrochloride Eye Ointment was used to prevent bacterial contamination after injection. Two replications were done. All pupae were kept at 25 °C until eclosion.

Within 48 hr after eclosion, virgin male moths were tested. One male was placed into a glass cultivate vessel (internal diameter = 180 mm) first and then one virgin female moth was added at the longest distance away from the male within the glass cultivate vessel. The movements of all male moths were recorded for 20 min and traces were analyzed by Tracker (http://www.cabrillo.edu/~dbrown/tracker). The results were graphed and analyzed in R. The relative expression of BmorSNMP1 and BmorOR1 in male antennae after RNAi was analysed by qRT-PCR using SYBR^^ Premix Ex Taq^TM^ (Perfect Real Time, Takara) according to the instructions manual for ABI 7500 fast real-time PCR system. The relative gene expression data were analyzed using the method described previoulsy ^26^. Data were further analysed using two-tailed Student’s t-test and graphed with GraphPad Prism 5.

### Split-ubiquitin yeast hybridization

Split-ubiquitin yeast hybridization system was performed using DUALmembrane pairwise interaction kit according to the manual (Dualsystems Biotech, Switzerland) and with the method used ^27^. Primers and constructs for fusion protein expression constructs were listed in Supplementary Table 4. Vectors were chosen to adapt for the transmembrane topology of proteins. In this study, pPR3N prey vector providing NubG and pBT3STE bait vector providing Cub were used in purpose. *SNMP1s* and *BmorOR2* were cloned into pPR3N prey, while cis-vaccinyl acetate receptor DmelOR22a and pheromone receptors (*BmorOR1*, *DmelOR67d* and *HarmOR13*) were cloned into pBT3STE bait vectors. *Saccharomyces cerevisiae* yeast (NMY32) were suspended amplified in YPDA (1% yeast extract, 2% Trytone, 2% Glucose and 0.02% Adenine) at 30ºC and 225rpm until OD600>1.5. Binary constructs were subsequently co-transformed into yeast which were then cultured in SD-Trp-Leu (SD-T/L) and SD-TrpLeu-His-Ade (SD-T/L/H/A) defective selection solid medium. The expression of three reporter genes of HIS3, ADE2 and LacZ in yeast could be induced due to the interaction between two fusion expressed proteins. The results are represented as positive clones in selection medium. The activity of β-galactosidase encoded by LacZ gene in yeast could be stained by X-gal solution at room temperature and then detected by microplate reader. Data were analyzed with GraphPad Prism 5 using the two-tailed Student’s t-test for comparisons between the potential pairwise interaction and negative control (pPR3N/pBT3STE).

## Acknowledgments

We thank Dr. Richard Vogt, Dr. Stephen C. Trowell for comments and discussion. Dr. Richard D. Newcomb and Dr. Selene van der Poel for poviding constructs of DmelOR22a, DmelOR67d and DmelSNMP1; Dr. Ting-Cai Cheng for assisting microarray data analysis, Dr. Faisal Younus and Dr. Olivia Leitch for providing *Drosophila* strain and assisting with dissections. This work was supported by China Postdoctoral Science Foundation (2013M531930), partially funded by National Natural Science Foundation of China (31530071) and funding from the Transformational Biology Capability Platform (TCP1:Dissecting Adaptive Potential) of the Commonwealth Scientific and Industrial Research Organisation, Australia. Dr. Wei Xu is the recipient of an Australian Research Council Discovery Early Career Researcher Award (DECRA) (DE160100382).

## Author Contributions

Hui-Jie Zhang conceived, designed and investigated the experiments, analyzed the data and wrote the manuscript.

Wei Xu performed some data analysis, revised and edited the manuscript.

Quan-mei Chen and Le-Na Sun detect the expression of BmorSNMPs and part of behaviour experiments on *Bombyx mori*.

Qing-You Xia co-ordinated the insect bioassay research, supervised and co-designed the work, and drafted the manuscript.

Alexie Papanicolaou co-ordinated the genomic research, co-designed the work, supervised the analysis, data analysis, drafted and revised the manuscript.

Dr. Alisha Anderson co-supervised the insect bioassay research, funded the lab for *Drosophila* and *Helicoverpa armigera* experiments.

All authors contributed to the manuscript, read and approved the final version. All authors report no conflicts of interest.

## Legend for Supplementary Material

Supplementary Resource 1 Video of single pair of male-female behavioural assays

Supplementary Resource 2 Example for those males lost sexual behavioural response elicited by female after RNAi

Supplementary Resource 3 Video of single pair of male-female behavioural assays. Male moth went away though male and female were placed close to each other directly.

Supplementary Resource 4 Video of competition behavioural assay after BmorSNMP1 dsRNA injection

Supplementary Resource 5 Video of competition behavioural assay after BmorOR1 dsRNA injection

Supplementary Table 1: Curated SNMPs within the Arthropoda

Supplementary Table 2: Other known SNMPs used in the phylogenetic analysis

Supplementary Table 3 Primers used in semi-qRT-PCR are listed as below

Supplementary Table 4 Primers used for split-ubiquitin yeast hybridization are listed as below.

Supplementary TABLE 5 The data of β-GAL experiments for pairwise interaction of proteins

## References

1 Vogt, R. G. et al. The insect SNMP gene family. Insect Biochem Mol Biol 39, 448–456, doi:10.1016/j.ibmb.2009.03.007 (2009).

2 Rogers, M. E., Steinbrecht, R. A. & Vogt, R. G. Expression of SNMP-1 in olfactory neurons and sensilla of male and female antennae of the silkmoth Antheraea polyphemus. Cell Tissue Res 303, 433–446 (2001).

3 Rogers, M. E., Krieger, J. & Vogt, R. G. Antennal SNMPs (sensory neuron membrane proteins) of Lepidoptera define a unique family of invertebrate CD36-like proteins. J Neurobiol 49, 47–61 (2001).

4 Rogers, M. E., Sun, M., Lerner, M. R. & Vogt, R. G. Snmp-1, a novel membrane protein of olfactory neurons of the silk moth Antheraea polyphemus with homology to the CD36 family of membrane proteins. J Biol Chem 272, 14792–14799 (1997).

5 Benton, R., Vannice, K. S. & Vosshall, L. B. An essential role for a CD36-related receptor in pheromone detection in Drosophila. Nature 450, 289–293, doi:10.1038/nature06328 (2007).

6 German, P. F., van der Poel, S., Carraher, C., Kralicek, A. V. & Newcomb, R. D. Insights into subunit interactions within the insect olfactory receptor complex using FRET. Insect Biochemistry and Molecular Biology 43, 138–145, doi:10.1016/j.ibmb.2012.11.002 (2013).

7 Forstner, M. et al. Differential expression of SNMP-1 and SNMP-2 proteins in pheromone-sensitive hairs of moths. Chem Senses 33, 291–299, doi:10.1093/chemse/bjm087 (2008).

8 Zhang, Y. N. et al. Differential Expression Patterns in Chemosensory and Non-Chemosensory Tissues of Putative Chemosensory Genes Identified by Transcriptome Analysis of Insect Pest the Purple Stem Borer Sesamia inferens (Walker). Plos One 8, doi:ARTN e69715 10.1371/journal.pone.0069715 (2013).

9 Zhang, J., Liu, Y., Walker, W. B., Dong, S. L. & Wang, G. R. Identification and localization of two sensory neuron membrane proteins from Spodoptera litura (Lepidoptera: Noctuidae). Insect Sci 22, 399–408, doi:10.1111/1744-7917.12131 (2015).

10 Liu, C. C., Zhang, J., Liu, Y., Wang, G. R. & Dong, S. L. Expression of Snmp1 and Snmp2 Genes in Antennal Sensilla of Spodoptera Exigua (Hubner). Arch Insect Biochem 85, 114–126, doi:10.1002/arch.21150 (2014).

11 Kirkness, E. F. et al. Genome sequences of the human body louse and its primary endosymbiont provide insights into the permanent parasitic lifestyle. Proceedings of the National Academy of Sciences of the United States of America 107, 12168–12173, doi:10.1073/pnas.1003379107 (2010).

12 Pearce, S. L. et al. Genomic innovations, transcriptional plasticity and gene loss underlying the evolution and divergence of two highly polyphagous and invasive Helicoverpa pest species. BMC Biol 15, 63, doi:10.1186/s12915-017-0402-6 (2017).

13 Papanicolaou, A., Joron, M., Mcmillan, W. O., Blaxter, M. L. & Jiggins, C. D. Genomic tools and cDNA derived markers for butterflies. Molecular Ecology 14, 2883–2897, doi:10.1111/j.1365-294X.2005.02609.x (2005).

14 Waterhouse, R. M., Tegenfeldt, F., Li, J., Zdobnov, E. M. & Kriventseva, E. V. OrthoDB: a hierarchical catalog of animal, fungal and bacterial orthologs. Nucleic Acids Res 41, D358–D365, doi:10.1093/nar/gks1116 (2013).

15 Jin, X., Ha, T. S. & Smith, D. P. SNMP is a signaling component required for pheromone sensitivity in Drosophila. Proc Natl Acad Sci U S A 105, 10996–11001, doi:10.1073/pnas.0803309105 (2008).

16 Gomez-Diaz, C., Reina, J. H., Cambillau, C. & Benton, R. Ligands for pheromone-sensing neurons are not conformationally activated odorant binding proteins. PLoS Biol 11, e1001546, doi:10.1371/journal.pbio.1001546 (2013).

17 Liu, S. et al. Identification and Characterization of Two Sensory Neuron Membrane Proteins from Cnaphalocrocis Medinalis (Lepidoptera: Pyralidae). Arch Insect Biochem 82, 29–42, doi:10.1002/arch.21069 (2013).

18 Liu, S. et al. Molecular Characterization of Two Sensory Neuron Membrane Proteins From Chilo suppressalis (Lepidoptera: Pyralidae). Annals of the Entomological Society of America 106, 378–384, doi:10.1603/AN12099 (2013).

19 Gu, S. H. et al. Molecular identification and differential expression of sensory neuron membrane proteins in the antennae of the black cutworm moth Agrotis ipsilon. Journal of Insect Physiology 59, 430–443, doi:10.1016/j.jinsphys.2013.02.003 (2013).

20 Liu, Y., Gu, S., Zhang, Y., Guo, Y. & Wang, G. Candidate olfaction genes identified within the Helicoverpa armigera Antennal Transcriptome. Plos One 7, e48260, doi:10.1371/journal.pone.0048260 (2012).

21 Li, P. Y. & Qin, Y. C. Molecular cloning and characterization of sensory neuron membrane protein and expression pattern analysis in the diamondback moth, Plutella xylostella (Lepidoptera: Plutellidae). Applied Entomology and Zoology 46, 497–504, doi:10.1007/s13355-011-0067-5 (2011).

22 Nichols, Z. & Vogt, R. G. The SNMP/CD36 gene family in Diptera, Hymenoptera and Coleoptera: Drosophila melanogaster, D. pseudoobscura, Anopheles gambiae, Aedes aegypti, Apis mellifera, and Tribolium castaneum. Insect Biochem Mol Biol 38, 398–415, doi:10.1016/j.ibmb.2007.11.003 (2008).

23 Altschul, S. F. et al. Gapped BLAST and PSI-BLAST: a new generation of protein database search programs. Nucleic Acids Res 25, 3389–3402, doi:DOI 10.1093/nar/25.17.3389 (1997).

24 Edgar, R. C. MUSCLE: a multiple sequence alignment method with reduced time and space complexity. Bmc Bioinformatics 5, 1–19, doi:Artn 113 10.1186/1471-2105-5-113 (2004).

25 Stamatakis, A., Ludwig, T. & Meier, H. RAxML-III: a fast program for maximum likelihood-based inference of large phylogenetic trees. Bioinformatics 21, 456–463, doi:10.1093/bioinformatics/bti191 (2005).

26 Livak, K. J. & Schmittgen, T. D. Analysis of relative gene expression data using real-time quantitative PCR and the 2(T)(-Delta Delta C) method. Methods 25, 402–408, doi:10.1006/meth.2001.1262 (2001).

27 Lentze, N. & Auerbach, D. Membrane-based yeast two-hybrid system to detect protein interactions. Curr Protoc Protein Sci **Chapter 19**, Unit 19 17, doi:10.1002/0471140864.ps1917s52 (2008).

